## Supplementary figures and images for "Psychological stress disturbs bone metabolism via miR-335-3p/Fos signaling in osteoclast"

### Figure3 - figure supplement 1

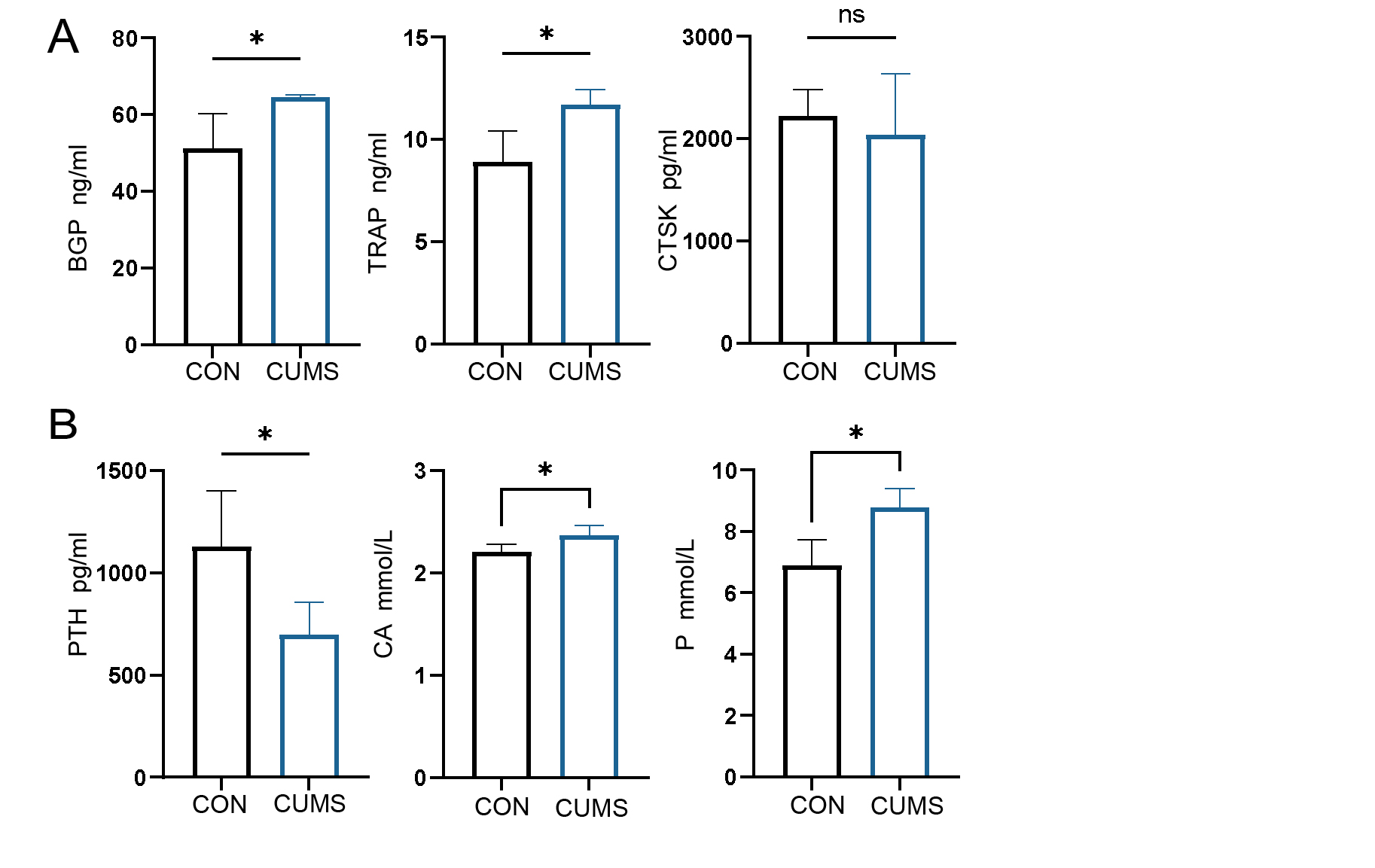

### Figure3 - figure supplement 2

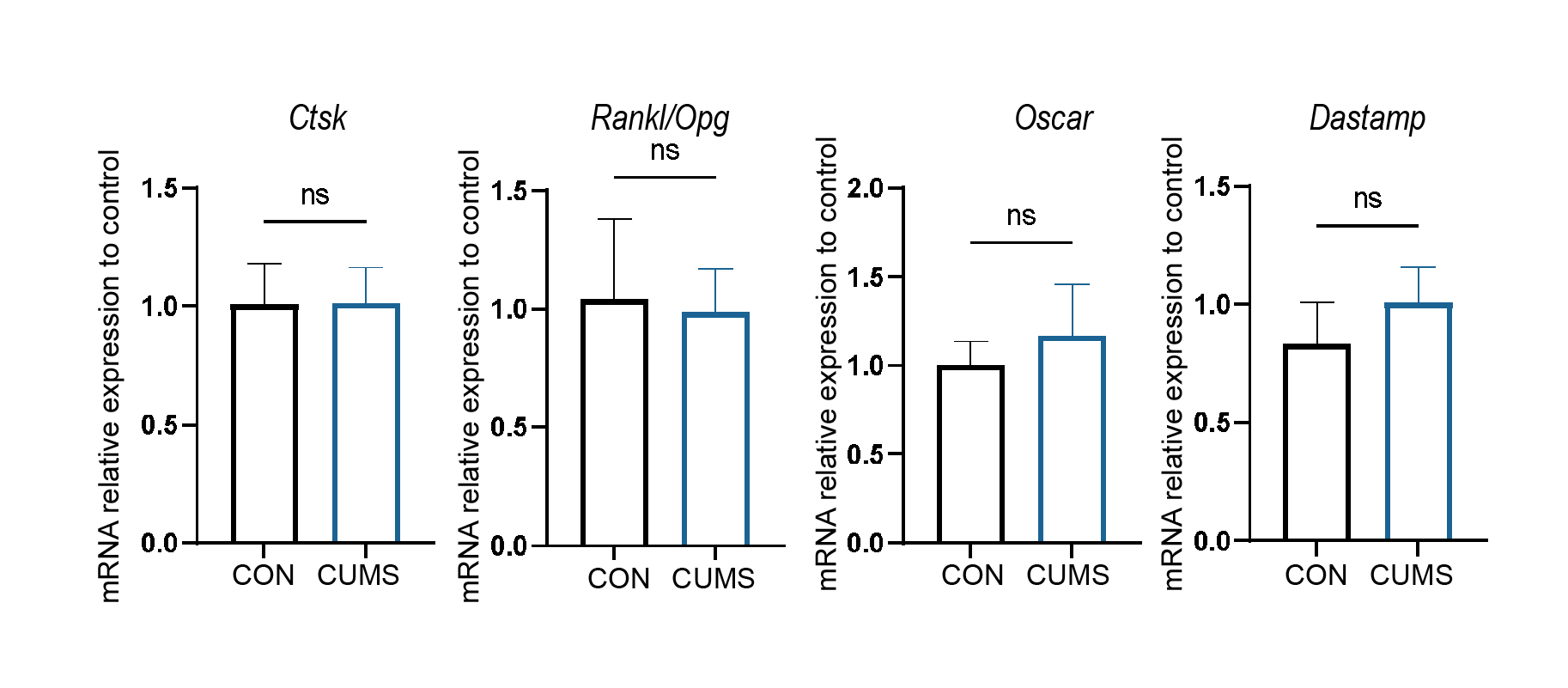

### Figure4 - figure supplement 1

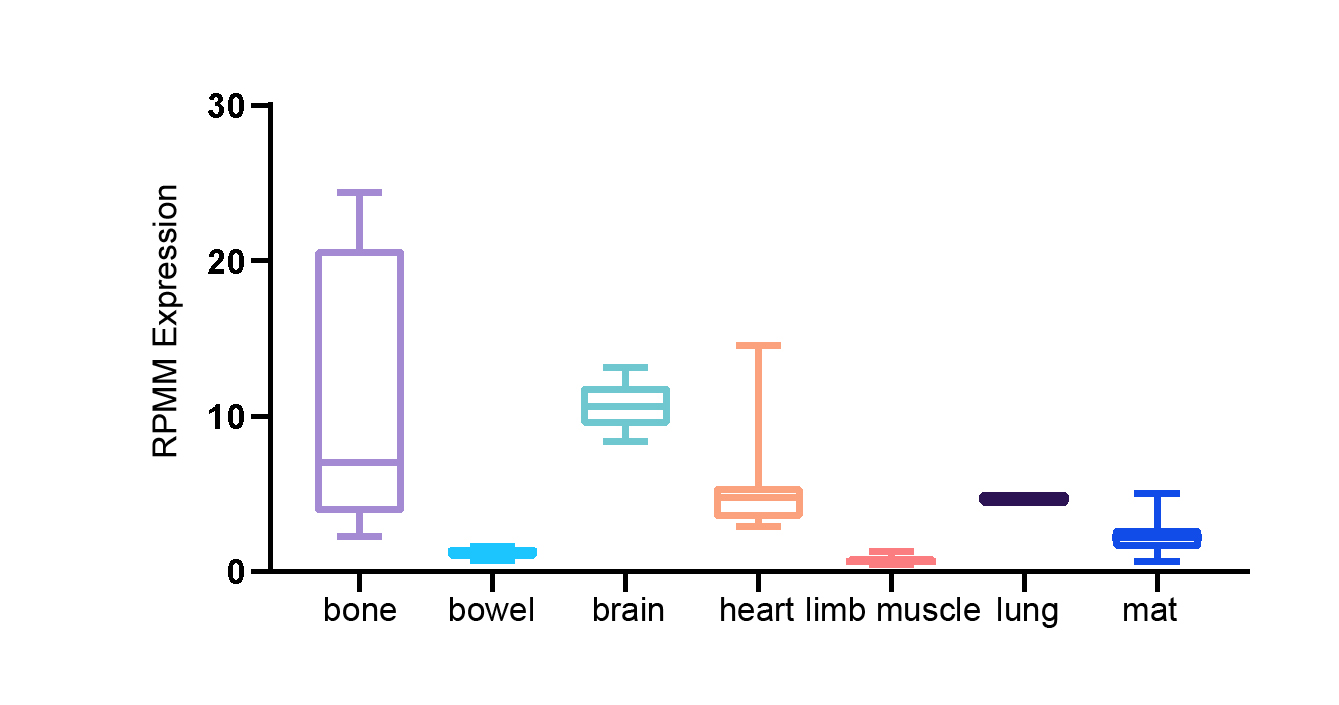

### Figure5 - figure supplement 1

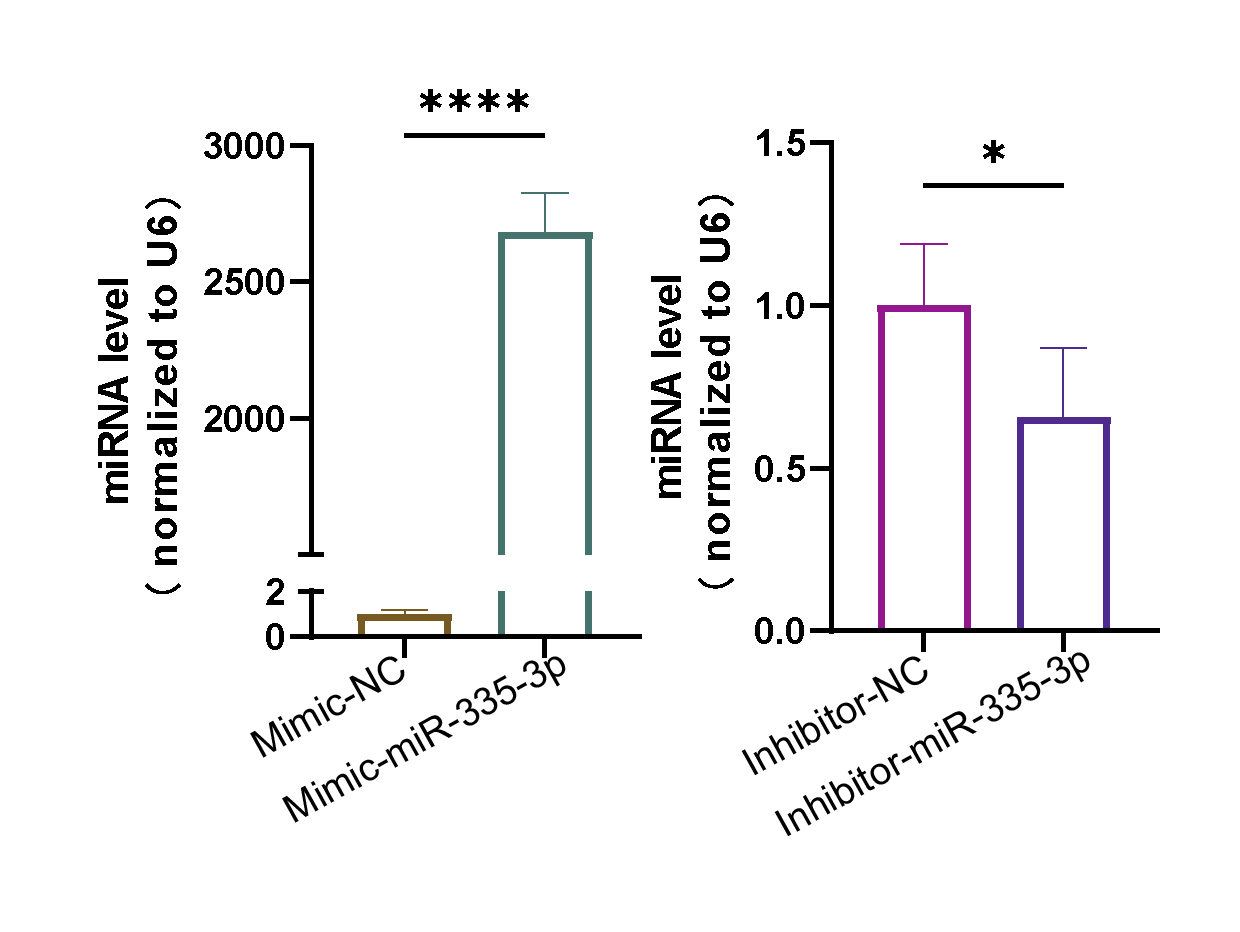

### Figure6 - figure supplement 1

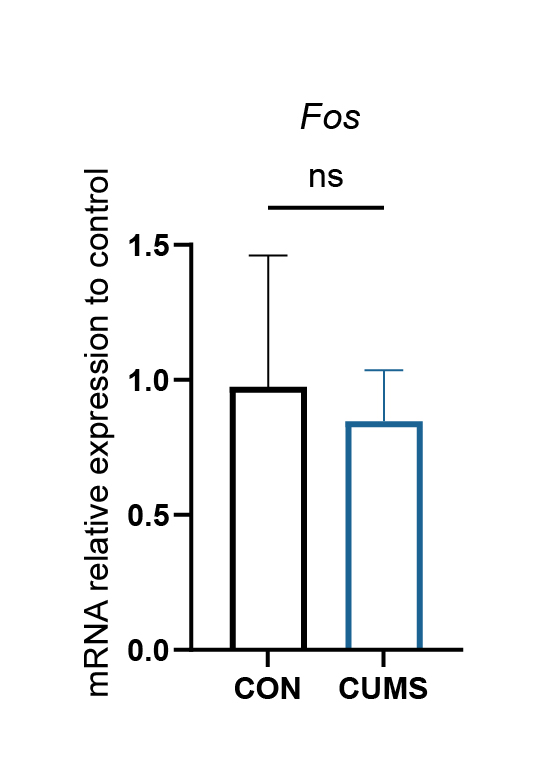
